# Isolation, identification, and characterisation of the malachite green detoxifying bacterial strain *Bacillus pacificus* ROC1 and the azoreductase AzrC

**DOI:** 10.1101/2024.08.29.610417

**Authors:** Shanza Bibi, Callum W. Breeze, Vusqa Jadoon, Anum Fareed, Alina Syed, Rebecca L. Frkic, Habiba Zaffar, Muhammad Ali, Iftikhar Zeb, Colin J. Jackson, Tatheer Alam Naqvi

## Abstract

Malachite green (MG) is used as a dye for materials such as wood, cotton, and nylon, and is used in aquaculture to prevent fungal and protozoan diseases. However, it is highly toxic, with carcinogenic, mutagenic, and teratogenic properties, resulting in bans worldwide. Despite this, MG is still frequently used in many countries due to its efficacy and economy. MG is persistent in the environment and so requires degradative intervention. In this work we isolated *Bacillus pacificus* ROC1 strain from a salt flat in Pakistan that had the ability to aerobically detoxify MG, as determined by bacterio- and phyto-toxicity assays. We demonstrate immobilized *B. pacificus* ROC1 can effectively detoxify MG, which highlights a potential method for its biodegradation. Genomic sequencing identified three candidate azo-reductases within *B. pacificus* ROC1 that could be responsible for the MG-degrading activity. These were cloned, expressed and purified from *Escherichia coli*, with one (AzrC), catalyzing the reduction of MG to leuco-MG *in vitro.* AzrC was crystallised and MG was captured within the active site in a Michaelis complex, providing structural insight into the reduction mechanism. Altogether, this work identifies a bacterium capable of aerobically degrading a major industrial pollutant and characterizes the molecular basis for this activity.

## Introduction

Malachite green (MG), a triphenylmethane dye, is a water-soluble dye that is often used to dye materials such as wood, leather, nylon, and cotton (Srivastava et al. 2004). MG is also used in aquaculture as it prevents diseases such as saprolegniasis caused by the mould *Saprolegnia*, and white spot disease caused by the protozoan *Ichthyophthirius multifiliis* (Bruno et al. 2011; Scholz 1999; West 2006). Despite the efficacy of MG, it has been banned in many nations, including China, the EU, and the US, due to negative health effects (EC 2004; Liu et al. 2012; Srivastava et al. 2004). MG can accumulate in organisms, along with the lipophilic reduced leuco form (LMG), and has been documented to be carcinogenic, teratogenic, and mutagenic (Allen and Hunn 1986; Andersen et al. 2006; Culp et al. 2006; Fessard et al. 1999; Gavrilenko et al. 2019; Srivastava et al. 2004; Stammati et al. 2005; Vilhena et al. 2023; WHO 2008; 2009). Low levels of MG (0.1 mg/L) have been shown to be toxic to mammalian cells, lower serum calcium levels, and reduce fertility (Chaturvedi et al. 2013; Clemmensen et al. 1984; Srivastava et al. 1995). MG, being a chromophore in visible wavelengths, also has environmental impacts; it decreases the transmission of light through waterbodies, resulting in decreased photosynthetic activity, and thus oxygen (Nelson and Hites 1980). Yet, MG is still used in some developing nations and is used illegally in others, due to its efficacy and cost efficiency; in the dyeing process 15% of dye is lost directly to wastewater (Bilandzic et al. 2012; Carmen and Daniela 2012; Conti et al. 2015; Srivastava et al. 2004; WHO 2009). Indeed, despite bans on MG, eels in German and Belgian waterways were found to contain MG and LMG, and in 2022 and 2023 shrimp imports to Europe were found to contain MG and LMG (Belpaire et al. 2015; EC 2022; 2023; Schuetze et al. 2008).

The complex aromatic structure of MG makes it recalcitrant to degradation, resulting in accumulation in the ecosystem, but consistent with their intended use as a dye needing to be resistant to light, water, oils, and chemicals (Mnif et al. 2015). Chemical degradation methods, such as ozonation, the use of photo-Fenton reagents, microwave-assisted catalysis, sono-photolysis, and photoelectrocatalysis have been developed for the degradation of MG (Berberidou et al. 2007; Bouafia-Chergui et al. 2012; Kusvuran et al. 2011; Liu et al. 2014; Mishra et al. 2019; Venkatesh et al. 2017; Yang et al. 2010). However, chemical methods are held back by their cost, production of secondary pollutants, or low efficiency, which can result from salinity, pH, or temperature of wastewater.

A promising alternative to chemical degradation methods is biodegradation, using whole bacterial or fungal cells for complete detoxification of MG. This ability has been observed in multiple organisms, including *Saccharomyces cerevisiae*, *Streptomyces exfoliatus*, and *Bacillus vietnamensis* (Abu-Hussien et al. 2022; Jadhav and Govindwar 2006; Kabeer et al. 2019). Individual enzymes have also been identified as being able to degrade MG, such as the metalloenzymes laccase and manganese peroxidase, and the NADH dependent triphenyl methane reductase (TMR) (Kim et al. 2008; Xu et al. 2020; Yang et al. 2015; Yang et al. 2016). TMR has been described to reduce MG at the central C10 forming LMG, whereas laccase and manganese peroxidase have been identified as capable of oxidising MG to many smaller molecules. A *de novo* metalloenzyme has also been developed based off myoglobin for the degradation of MG (Xiang et al. 2021).

In this work, we isolated and characterised a bacterial strain from dye-contaminated salt flats in Pakistan, *Bacillus pacificus* ROC1 (ROC1). We identified ROC1 as being able to completely, and aerobically, decolourise MG. Additional experiments monitoring bacterio- and phyto-toxcity identified that ROC1 was not only able to decolourise, but also detoxify MG, suggesting a substantial breakdown of MG. We performed genomic sequencing of ROC1, alongside cell lysate assays, to identify an azoreductase capable of MG decolourisation. AzrC was recombinantly expressed and was observed to decolourise MG via reduction to LMG, as determined by LCMS analysis. The formation of LMG was supported by the solved AzrC-MG co-crystal structure, which captured MG and AzrC in a Michaelis complex.

## Results

### Malachite green degradation in bacterial culture and immobilized cells

To identify the ability of ROC1 to utilise MG as a sole carbon or nitrogen source, ROC1 was screened on different minimal media, containing either a carbon source, a nitrogen source, or neither, all with 50 ppm MG. ROC1 was unable to grow on any minimal media, indicating that ROC1 was unable to use MG as a sole carbon or nitrogen source. However, ROC1 grew well with LB media containing 250 ppm MG, and visibly decolourised the dye, indicating that ROC1 likely co-metabolises MG.

To quantify the ability of ROC1 to decolourise MG, ROC1 was incubated with various concentrations of MG (Figure 1A). The duration of treatment affected the overall decolourisation of MG and it took more time to decolourise fully when higher concentrations were used. For the 50 ppm MG culture, 100% decolorization was observed after 72 hours of treatment, while 100% decolorization for 100 to 500 ppm was observed after 96 hours of treatment and for 750 ppm decolourisation required 144 hours. However, in the presence of 1,000 ppm only 41% decolourisation was recorded after 144 hours or greater, a significant decrease compared to all other concentrations, suggesting MG could have a toxic effect on the cells at such high concentrations.

**Figure 1.**
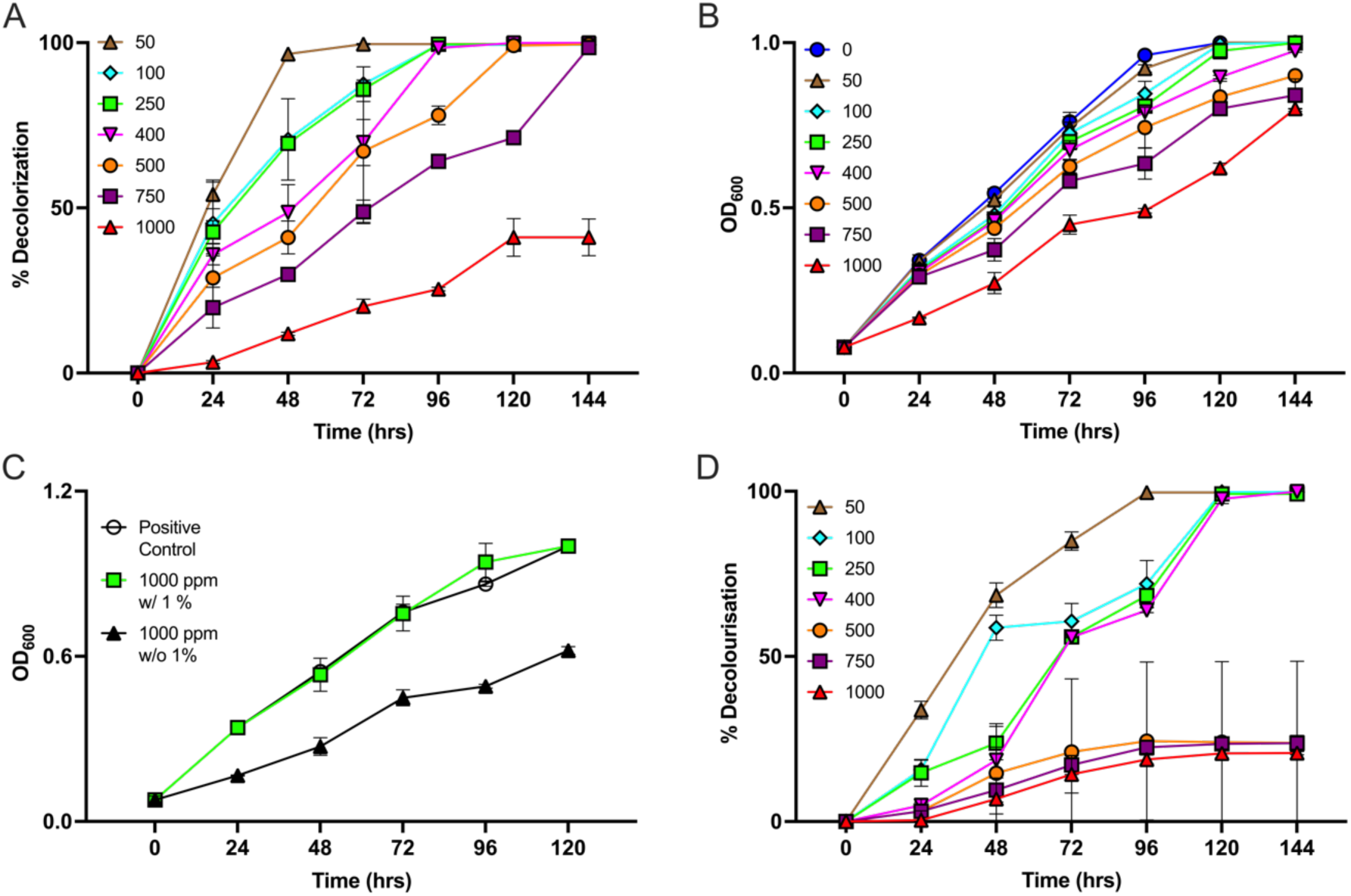
ROC1 decolourisation of MG. A – Decolourisation of 50-1,000 ppm MG measured at 620 nm over 144 hours by free ROC1 ±SD (N=2). B – Optical density measured at 600 nm of ROC1 cultures grown over 144 hours in the presence of 0-1,000 ppm MG ±SD (N=2). C – Optical density of free ROC1 measured at 600 nm in the addition of 1% (v/v) media with 0 ppm (white) and 1000 ppm (green) MG, and with 1000 ppm without media addition (black) ±SD (N=2). D – Decolourisation of 50-1,000 ppm MG measured at 620 nm over 144 hours by immobilised ROC1 ±SD (N=2). Data presented with significance in Supporting Information 3.

To investigate the potential inhibitory effect of high MG concentrations on cell growth, the growth of ROC1 was recorded along with dye decolourisation every 24 hours. In the presence of all concentrations of MG, the OD_600_ increased. However, it was found that faster growth was observed at lower concentrations up until 500 ppm (OD_600_ 1), and at higher concentrations of 750 and 1000 ppm the growth was significantly inhibited (Figure 1B). Similarly, when measuring colony forming units, wet biomass, and dry biomass (Supporting Information 1), the data indicated that all concentrations of MG allowed cell growth, but that growth was slower at higher MG concentrations, suggesting that MG does have an inhibitory effect against ROC1 growth, which becomes apparent at high concentrations.

To investigate approaches to prevent MG inhibition of ROC1 growth, LB media was adjusted with 0.5% (w/v) and 1% (w/v) glucose alongside 1,000 ppm MG. 1,000 ppm MG was chosen because this concentration was not fully decolourised and resulted in reduced ROC1 growth, permitting identification of improvement. No significant improvement in decolourisation was observed with the addition of glucose (Supporting Information 2). Alternatively, the continuous addition of 1% (v/v) fresh LB was conducted every 24 hours, resulting in 100% decolourisation of MG after 120 hours (Supporting Information 2). The addition of fresh LB also affected ROC1 growth positively, with an insignificant difference in OD_600_ after 120 hours between samples with 0 and 1,000 ppm MG, and a significant difference between 1,000 ppm samples with and without media addition (Figure 1C). These results suggest that the addition of fresh media increased the cell biomass, which had a positive impact on the decolourisation of MG at high concentrations.

Immobilised cells (IMC) are an attractive substitute for free cell biodegradation due to their reusability and stability (Kulkarni and Chaudhari 2007). IMC of ROC1, prepared using an agar entrapment method previously described (Fareed et al. 2022; Woodward 1988), were tested to determine their ability to decolourise MG. After 144 hours of treatment, 100% decolourisation was observed for concentrations of 50-400 ppm (Figure 1D). However, at 500-1,000 ppm MG approximately 20% decolourisation was recorded after 144 hours of treatment. This reduced decolourisation, as compared to the free cells, could be a result of reduced biomass.

### Toxicity of ROC1 products

Given that ROC1 was capable of MG decolorization, ROC1 could prove useful if it were able to degrade MG into non-toxic molecules. To test for bacteriotoxicity of MG degradation products, both Gram-positive and Gram-negative strains, *Staphylococcus aureus* and *Escherichia coli*, respectively, were used. Both strains exhibited growth inhibition when exposed to MG, with a positive relationship between clearance zone and MG concentration, demonstrating the toxicity of MG to bacteria (Figure 2A,B). Alternatively, MG treated with ROC1 did not inhibit growth of either bacteria.

**Figure 2.**
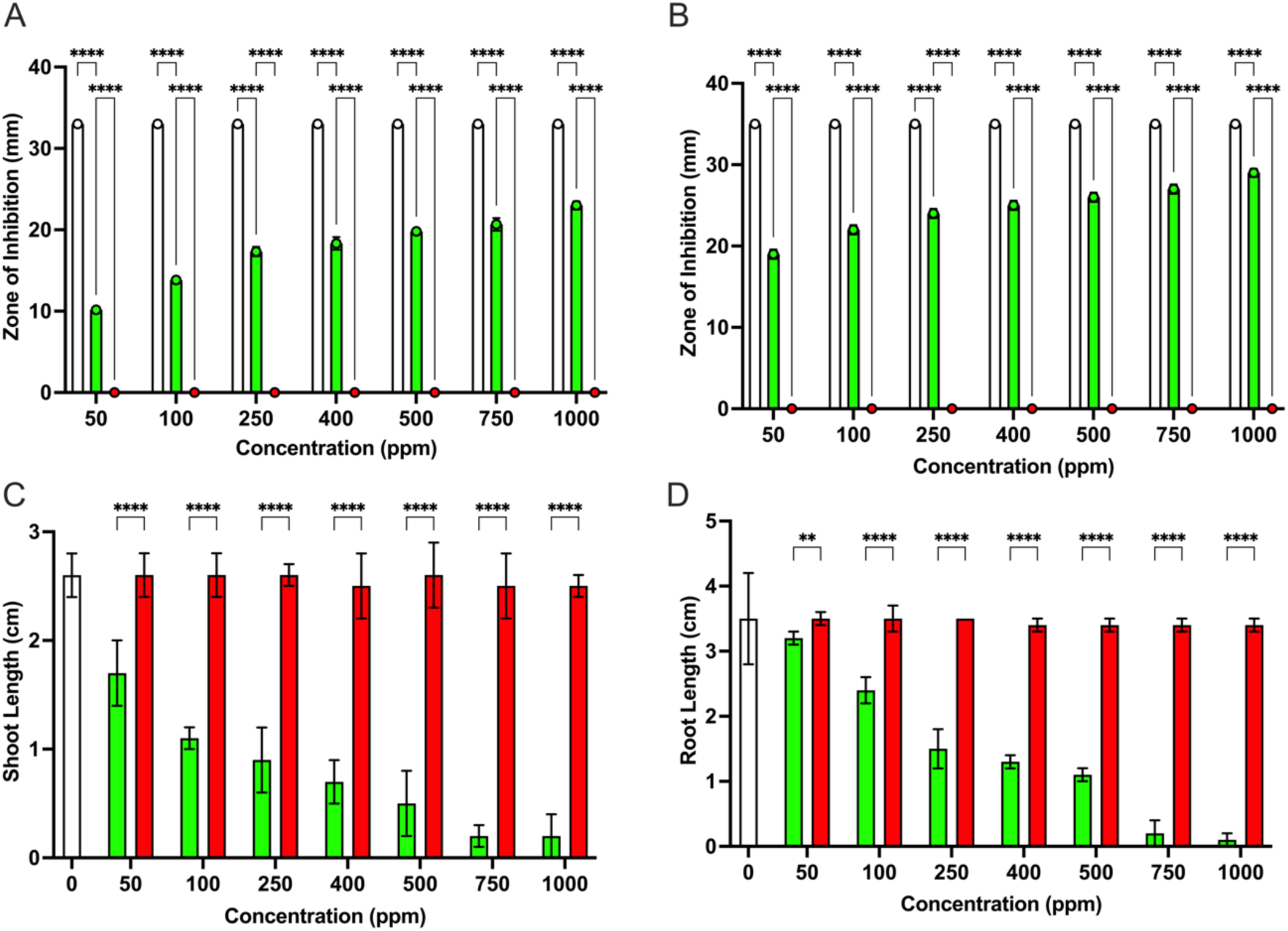
ROC1 treated MG toxicity. A – Clearance zones of *E. coli* when grown with kanamycin (white), untreated MG (green), and treated MG (red) ±SD. Asterisks indicate significance (N=3). B – Clearance zones of *S. aureus* when grown with kanamycin (white), untreated MG (green), and treated MG (red) ±SD. Asterisks indicate significance (N=3). C – Shoot length of *Solanum lycopersicum* when incubated with water (white), untreated MG (green), and treated MG (red) ±SD. Asterisks indicate significance (N=7). D – Root length of *Solanum lycopersicum* when incubated with water (white), untreated MG (green), and treated MG (red) ±SD. Asterisks indicate significance (N=7).

To test for phytotoxicity, the percentage of seed germination of the commercially important *Solanum lycopersicum* variety Riogrande (tomato) was assessed, along with the root and shoot length of germinated seeds. A substantial increase in percentage of seed germination was measured when seeds were incubated with water containing MG and treated with ROC1 *vs.* untreated water, with a negative relationship between percentage of seed germination and MG concentration (Supporting Information 4). At 50 ppm MG there was a 60% germination rate and 13% at 1,000 ppm. In ROC1 treated water, the germination rate was above 80% for all MG concentrations, indicating that the phytotoxic effect of MG is ameliorated by ROC1-mediated MG degradation. A similar negative relationship was observed for root and shoot length (Figure 2C, D): at 1,000 ppm MG the root and shoot length were negligible, whereas in ROC1 treated samples the root and shoot lengths were significantly longer, suggesting that any metabolites formed after ROC1-mediated degradation were not phytotoxic. Altogether, these results indicate that ROC1 can completely detoxify MG and that any metabolites produced after treatment by ROC1 were non-toxic to the bacteria and plants tested.

### Identifying the enzymatic basis for MG degradation

The cell free extract (CFE) of ROC1 was used to identify the enzyme involved in MG degradation, with specific testing for laccase and reductase activity since these enzyme families have previously been observed to catalyze MG degradation (Kim et al. 2008; Xu et al. 2020; Yang et al. 2015). The reductase assay revealed a gradual decrease in MG absorbance at 620 nm over time with CFE (Figure 3). When additional NADH was provided there was a significant increase in the rate of decolourisation. Alternatively, no laccase activity was detected in the ROC1 CFE. This suggested that the enzyme responsible for MG decolourisation was an NADH-dependent reductase.

**Figure 3.**
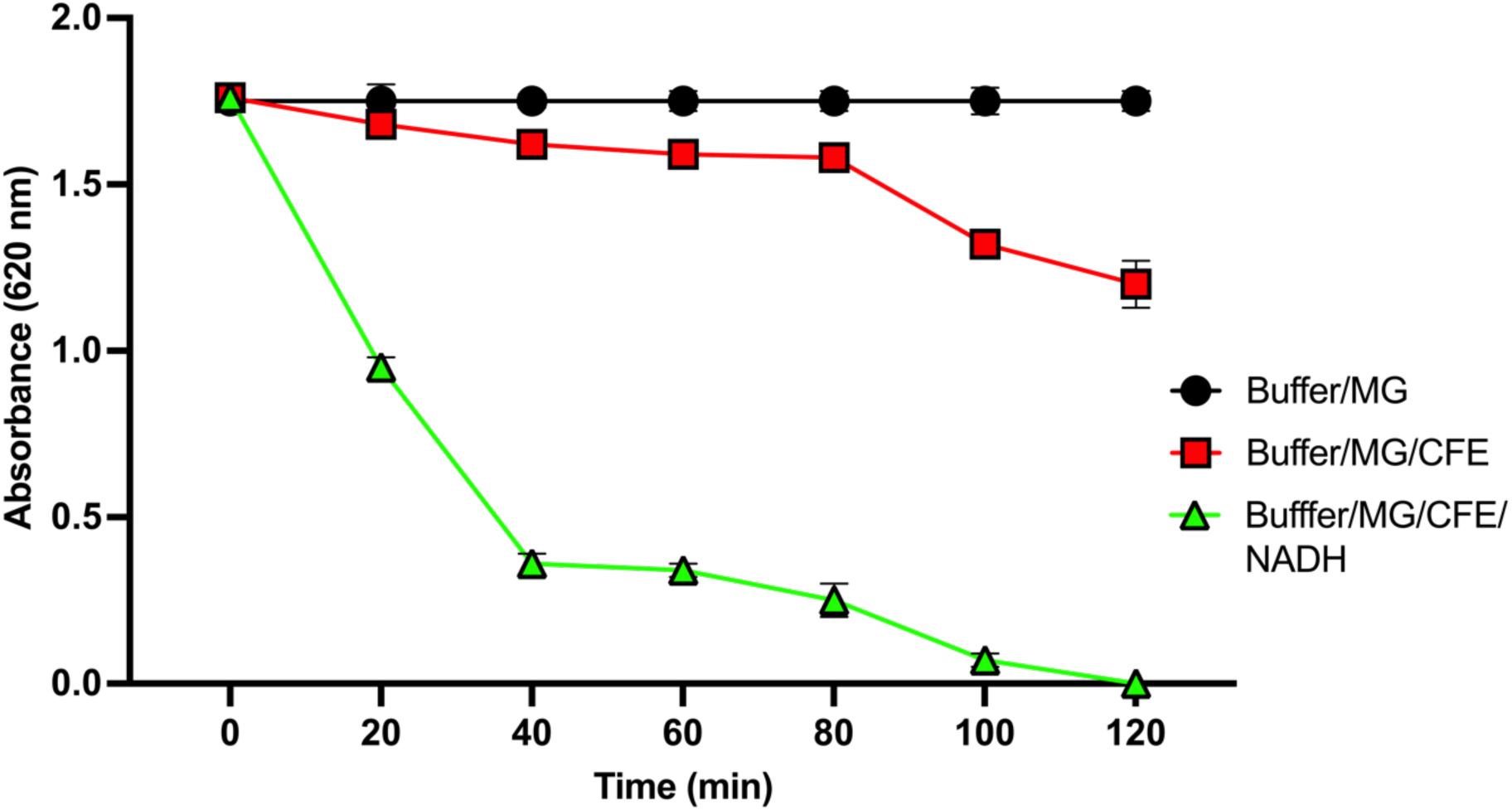
Reductase activity in the CFE. MG absorbance (620 nm) measured over two hours in buffer (black), buffer with CFE (red), and buffer with CFE and NADH (green) ±SD (N=3). Data presented with significance in Supporting Information 5.

To identify MG degrading enzymes, the ROC1 genome was sequenced with 98.82% completeness, and revealed a genome length of 5,189,869 bp (Table 1), with a GC content of 35.4%. The ROC1 genome has 5,384 predicted protein encoding genes, 102 predicted tRNA genes, and 32 predicted rRNA genes. The circular genome generated by Proksee (Supporting Information 6) displays the genome characteristics, including gene distribution and GC content (Grant et al. 2023). RAST annotation of the ROC1 genome categorised 5,384 genes into 27 subsystems (Figure 4A). A total of 23% of its genes are present in the largest subsystem, “amino acids and derivatives” with 326 members. “Metabolism of aromatic compounds” contained 12 predicted genes.

**Figure 4.**
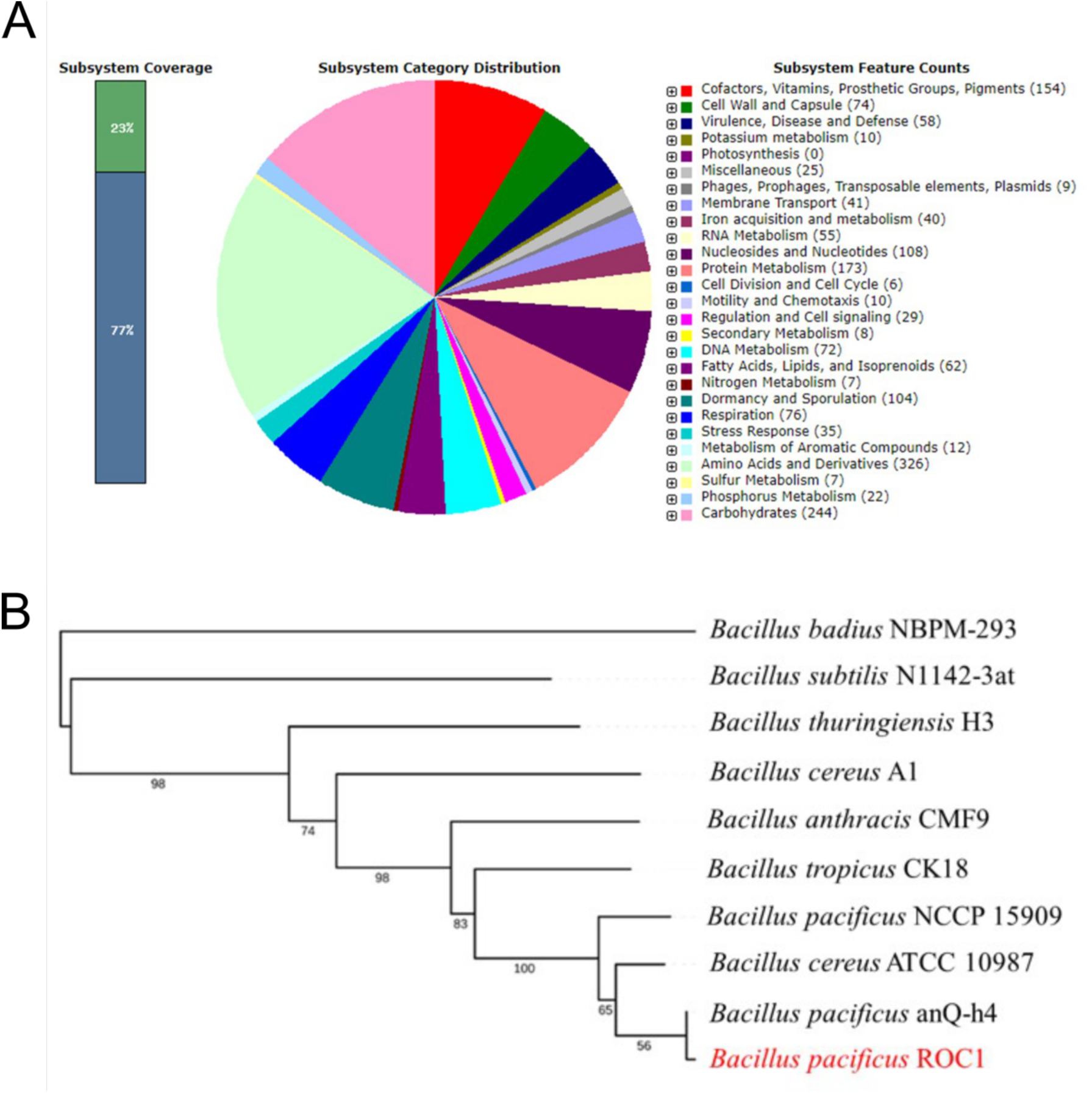
ROC1 genome. A – Functional annotation of ROC1 genomic genes using RAST. B – Phylogenetic tree of the relationship between ROC1 (red) and other *Bacillus* species. The branch numbers are GBDP pseudo-bootstrap values from 100 replicates.

**Table 1.** Statistics of the genome assembly. Statistics resulting from whole genome Illumina sequencing of *Bacillus pacificus* ROC1.

| Attribute | Value |
| --- | --- |
| No. of contigs | 33 |
| No. of scaffolds | 1 |
| Genome length (bp) | 5,189,869 |
| GC content (%) | 35.4 |
| Protein coding genes | 5,384 |
| rRNA | 32 |
| tRNA | 102 |

A phylogenetic tree (Figure 4B) was constructed using the Type Strain Genome Server (TYGS) for pairwise comparison between genomes using Genome Blast Distance Phylogeny (GBDP) and genetic distance calculator (Meier-Kolthoff et al. 2013; Meier-Kolthoff and Goker 2019). Both average nucleotide identity (ANI) and average amino acid identity (AAI) were used for phylogenetic analysis. ROC1 had the highest similarity with *B. pacificus* anQ_h4 with an ANI value of 100%. The lowest similarity was observed with *B. badius* NBPM_293 with an ANI value of 65.79%. Regarding AAI, ROC1 had the highest similarity with *B. pacificus* anQ_h4, *B. pacificus* NCCP_15909 and *Bacillus cereus* ATTC_10987 with AAI values of 100%. The lowest level of protein similarity was observed with *B. badius* NBPM_293 with an AAI value of 61.54%.

To identify candidate genes involved in MG degradation, the 12 genes categorised as as being involved in “metabolism of aromatic compounds” by RAST were examined. Of these initial 12, three were identified as putative MG degrading genes, via analysis using reference genes sourced from the Integrated Microbial Genomes (IMG) system, which were compared to the ROC1 genome (Markowitz et al. 2006). The products of these three genes were annotated as two FMN-dependent NADH-azoreductases (MDF0734648.1 and MDF0737692.1), AzrC and AzrA respectively, and one NADPH-dependent FMN reductase (MDF0734684.1) YhdA; His-tagged sequences, with Tobacco Etch Virus (TEV) cleave sites, are presented in Supporting Information 7. These three enzymes are found throughout the *Bacillus* genus and consist of 211, 208, and 178 residues, respectively. ROC1 AzrC and AzrA has sequence identities of 98% and 99% with AzrC (PDB-ID 3W79) and AzrA (PDB-ID 3W77) from *Bacillus* sp. B29, respectively, which have previously been shown to be capable of reducing numerous azo-dyes, including Methyl-Red, Orange I, and Acid Red 88 (Ooi et al. 2009; Ooi et al. 2007; Yu et al. 2014). ROC1 AzrC also had 57% sequence identity with Indigo reductase (PDB-ID 6JXN) from *Bacillus smithii* which could reduce indigo carmine to leuco-indigo carmine, along with reducing multiple azo-dyes (Yoneda et al. 2020). Whereas ROC1 YhdA had just 59% sequence identity with YhdA (PDB-ID 3GFR) from *Bacillus subtilis*, which has been demonstrated to reduce quinones and azo-dyes (Binter et al. 2009; Deller et al. 2006).

Purification of the azo-reductases was achieved by Ni(II)-affinity chromatography for initial assays (Supporting Information 8A). Elution fractions from both techniques were orange in colour, indicating the presence of a riboflavin based cofactor, and the SDS-PAGE contained a strong bands at ∼20-25 kDa for each protein, consistent with the calculated masses of 21.6, 25.2, and 25.0 kDa, for YhdA, AzrC, and AzrA, respectively. To determine which enzyme was responsible for the degradation of MG, all three enzymes were separately incubated with MG. Only AzrC was capable to decolourising MG and was capable of decolourising MG using both NADH and NADPH (Supporting Information 9). AzrC was then further purified suing size exclusion chromatography (Supporting Information 8B). Analysis of the 3D gel in the ImageLab software indicated that AzrC was >99% pure, with few impurities, indicating that AzrC was easily purified to a high extent. Both NADH and NADPH were used to determine Michaelis-Menten kinetics for AzrC with MG (Figure 5A and B). The *k*_cat_ was determined to be 0.942 s^-1^ and 0.611 s^-1^ for NADH and NADPH, respectively. The *K*_M_ was 935.2 μM (95% CI 794.3-1111) and 475.5 μM (95% CI 381.2-598.8) for NADH and NADPH, respectively.

**Figure 5.**
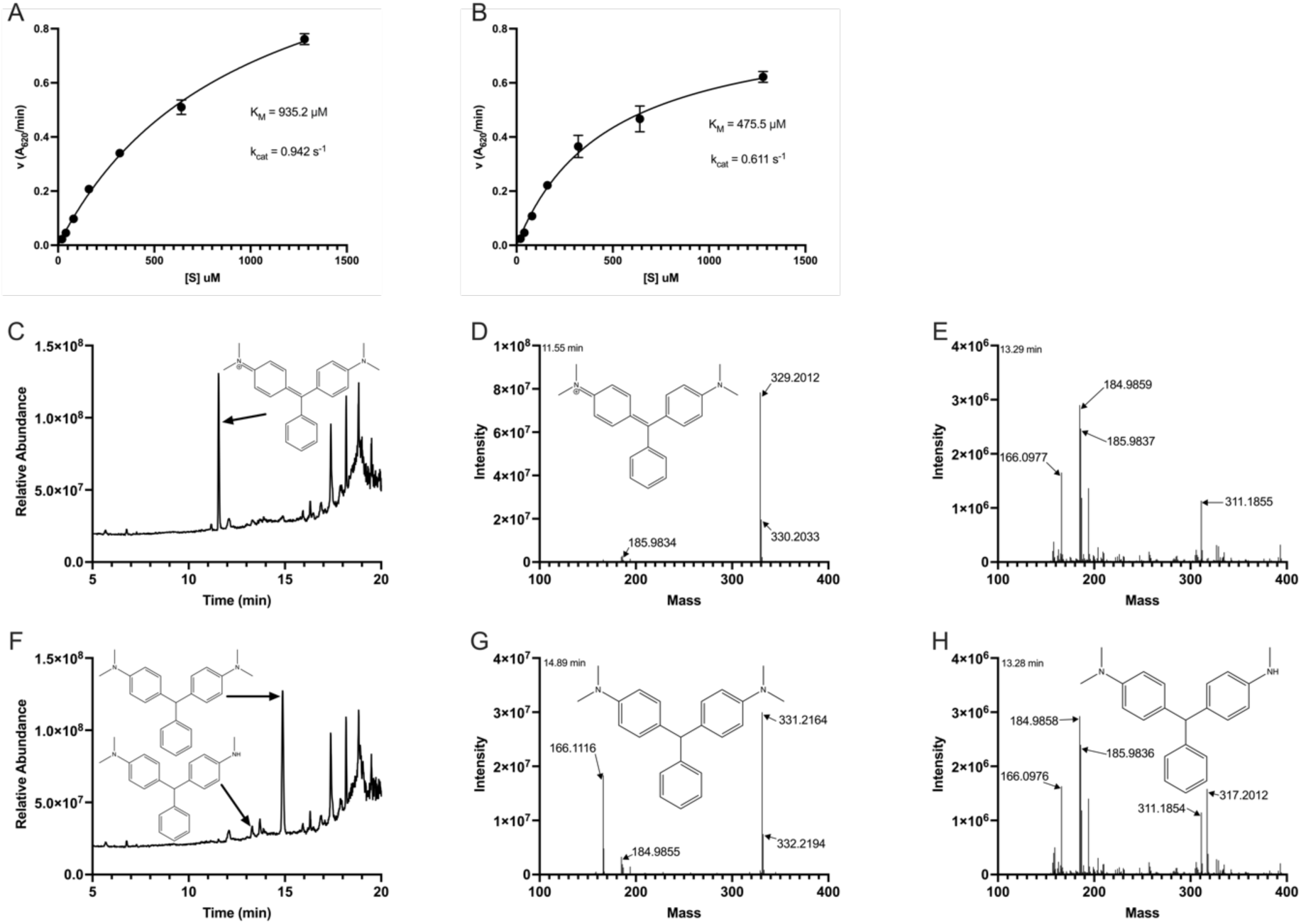
AzrC Reactivity. A – Michaelis-Menten plot of AzrC activity with NADH as a reducing agent. B – Michaelis-Menten plot of AzrC activity with NADPH as a reducing agent. C – LC chromatogram of the AzrC reaction with no reducing agent. D – Mass spectrum of MG at 11.55 minutes of the AzrC reaction with no reducing agent. E – Mass spectrum at 13.29 minutes of the AzrC reaction with no reducing agent. F – LC chromatogram of the AzrC reaction with reducing agent. G – Mass spectrum of LMG at 14.89 minutes of the AzrC reaction with reducing agent. H – Mass spectrum of desmethyl-LMG at 13.28 minutes of the AzrC reaction with reducing agent.

LCMS analysis was undertaken to determine reaction products of AzrC. The reaction with no NADH displayed a strong peak at 11.5 minutes on the LC, which was determined to be MG with mass 329.2012 (Figure 5C and D). Alternatively, the reaction with NADH displayed a peak, not at 11.5 minutes, but at 14.89 minutes, which was determined to be LMG with a protonated mass of 331.2164 (Figure 5F and G). A small peak at 13.28 minutes was also evident in the LC trace; this was identified to have a mass of 317.2012, consistent with desmethyl-LMG (Figure 5F and H). Desmethyl-LMG was not identified in the cofactor-free control reaction (Figure 5E). We are unsure of the mechanism of desmethyl-LMG formation, but due to over one minute difference in retention time compared to LMG, it appears that desmethyl-LMG was formed in the reaction.

### Structural characterization of AzrC-catalyzed MG reduction

Purified AzrC was co-crystallized with MG and the structure of the complex was solved to 2.35 Å and was deposited to the protein data bank (PDB) under PDB-ID 9C0W (Table 2). Despite a strong CC_1/2_ data were truncated to 2.35 Å, due to the presence of an ice ring. AzrC formed a dimer, with the FMN binding sites being located at the dimer interface. MG was bound at only one active site, due to the other being occupied by the N-terminus of a symmetry mate (Supporting Information 10). MG was modelled into a Polder electron density difference map (Figure 6A), showing that it was bound within the AzrC active site, with the cationic imine 3.7 Å from N5 of FMN (Figure 6B). Despite MG being bound within the predominantly hydrophobic and aromatic active site, there were few direct interactions as determined by Flatland Ligand Environment View (FLEV) (Coot) and PoseView (ProteinPlus), with FLEV indicating a π-cation interaction between F172 and the cationic imine of MG, and PoseView indicating π-π interactions between MG and Y151 and Y156 (Figure 6C and D), suggesting that AzrC likely has promiscuous substrate specificity. The π-cation interaction from F172 likely stands to aid in positioning the cationic imine proximal to FMN, suggesting reduction would occur at the cationic imine, with protonation at the central C10 forming LMG (Figure 6E).

**Figure 6.**
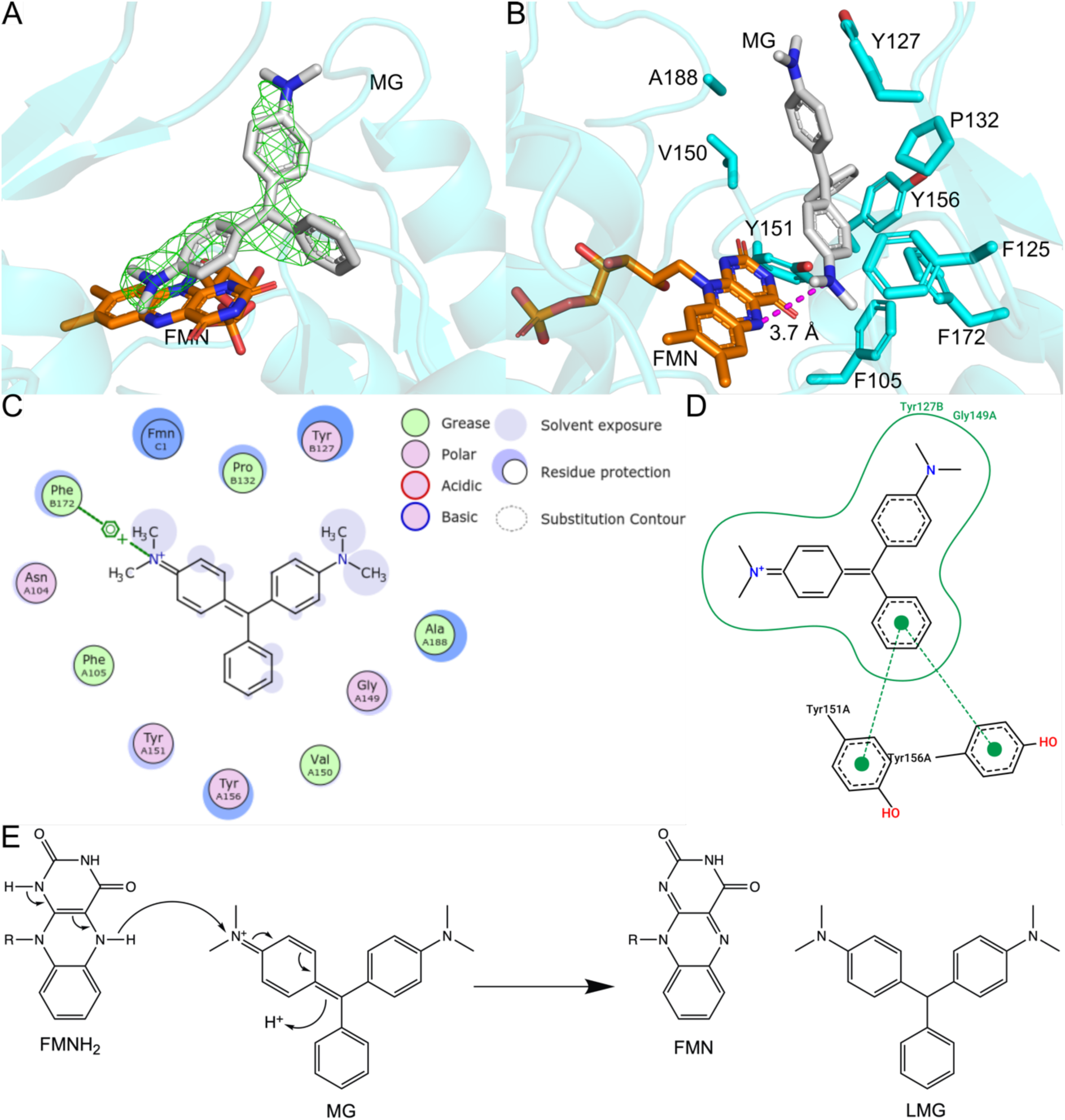
AzrC-MG co-crystal structure. Co-crystal structure of MG-bound AzrC (PDB 9C0W). A – Polder map at 5σ (green) of MG (grey). B – MG (grey) and surrounding residues of AzrC (cyan). C – Flatland ligand environment view (FLEV) of MG in 9C0W. D – Two-dimensional PoseView of MG in 9C0W. E – Proposed mechanism of MG reduction to LMG by AzrC.

**Table 2.**
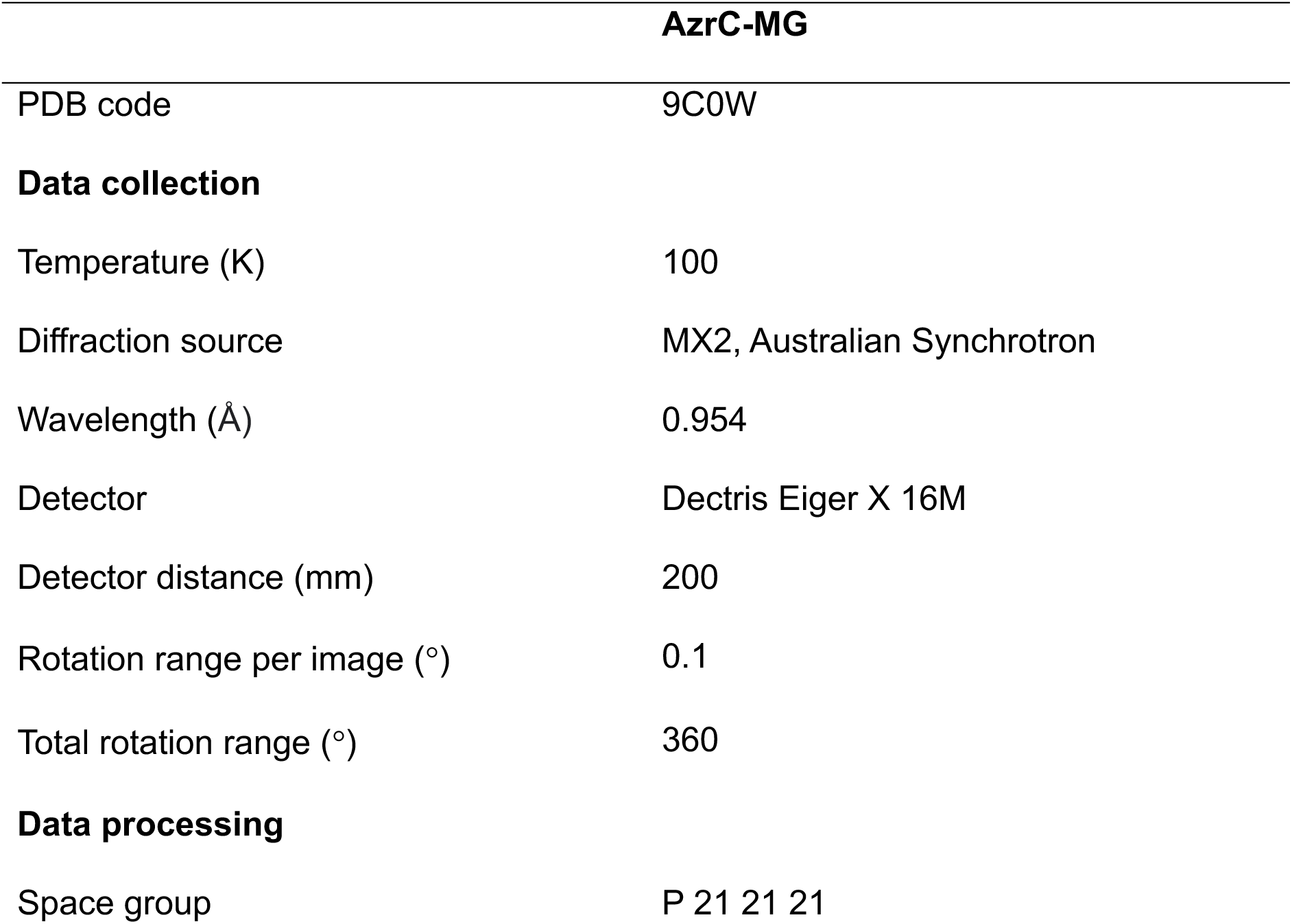
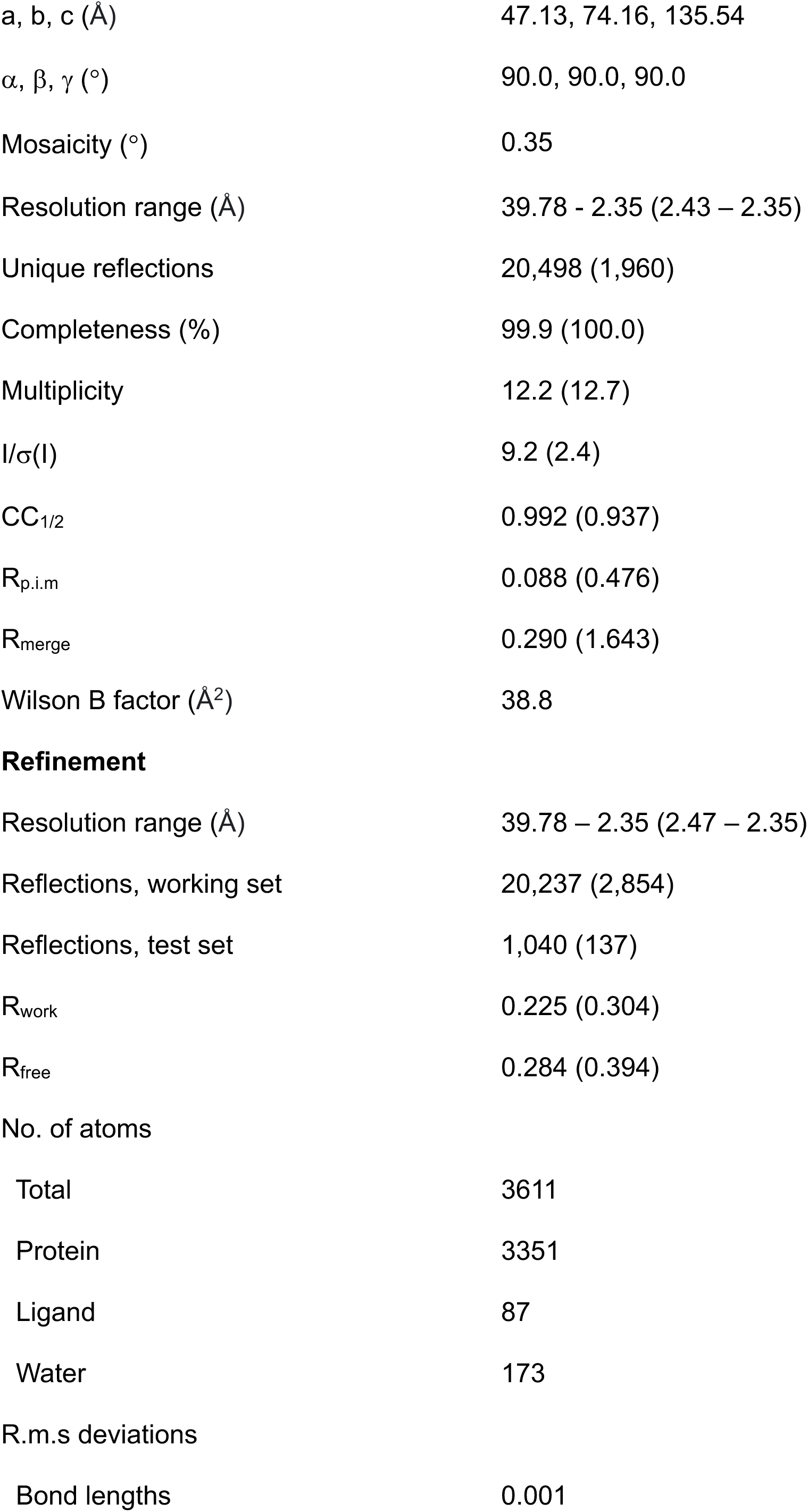
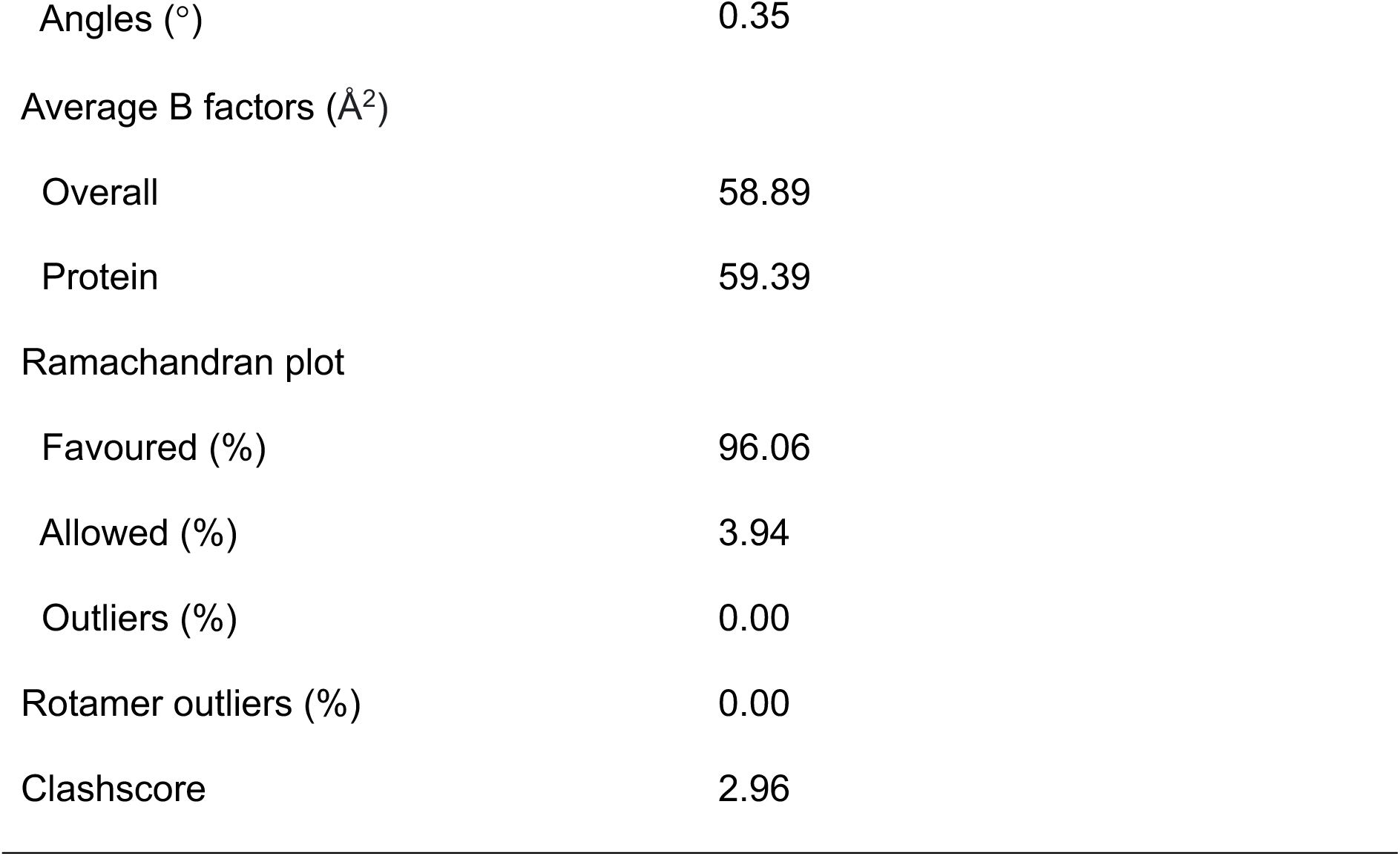
Crystallographic statistics. Data collection, processing, and refinement statistics for the crystal structure deposited as PDB entry 9C0W. Values in brackets are for the highest resolution shell.

## Discussion

In this work, we utilised a bioprospecting approach, and isolated *Bacillus pacificus* ROC1 from a MG contaminated site in the Kohat salt range in Pakistan. ROC1 was capable of aerobically degrading MG, and was able to detoxify MG, with the resulting treated water non-toxic to *E. coli*, *S. aureus*, and *S. lycopersicum* variety Riogrande. Despite the ability of ROC1 to degrade MG, it was unable to completely mineralise MG, suggesting that ROC1 works in consortia. The degradation and detoxification of MG suggest that ROC1 could have uses in MG biodegradation processes, due to the aerobic activity and halophilic nature of ROC1.

*Bacillus* species have been documented to be capable of aerobically degrading dyes: *B. subtilis* was documented to degrade orange II; *B. pseudomycoides* was documented to degrade methylene green, acid blue, and basic violet; and *B. cereus* was documented to degrade reactive orange 16 and reactive black 5 (Fareed et al. 2022; Hossain et al. 2022; Ikram et al. 2022). Whereas, for MG, most identified species have been of other genera, such as *Pseudomonas veronii*, *Streptomyces exfoliatus*, *Citrobacter sedlakii*, and *Saccharomyces cerevisiae* (Abu-Hussien et al. 2022; Jadhav and Govindwar 2006; Mnif et al. 2015; Song et al. 2020). There has been one description of a *Bacillus* genus member, *Bacillus vietnamensis* sp. MSB17, which was capable of MG degradation and detoxification. In the work investigating *B. vietnamensis*, there was an increase in tyrosinase, laccase, and manganese peroxidase activity during MG degradation, leading to the authors concluding the involvement of these enzymes in the process. However, there was no thorough investigation into these enzymes and there was no analysis of reductase activity (Kabeer et al. 2019).

In our work we investigated three enzymes from ROC1 for their ability to decolourise MG: AzrA, AzrC, and YhdA. Just one of these enzymes, AzrC, was identified as having activity with MG. AzrC from ROC1 was 98% identical to AzrC from *Bacillus* sp. B29 and 57% identical to Indigo reductase from *Bacillus smithii*. AzrC from *Bacillus* sp. B29 was documented to reduce methyl red, 1-(2-pyridylazo)-2-naphthol, acid red 88, and orange I, whereas indigo reductase from *B. smithii* was documented to reduce indigo carmine, methyl red, and acid red 88 (Ooi et al. 2009; Yoneda et al. 2020). However, neither has been shown to degrade MG. These dyes have similar properties to MG, and are all aromatic molecules, with hydrophobic regions, excluding indigo carmine which was used as a hydrophilic substitute for indigo. Also, the proposed mechanism for AzrC from *B*. sp. B29 reduction of orange I and acid red 88 involved reduction at an electrophilic carbonyl carbon ∼3.5 Å from N5 of FMN (Yu et al. 2014). The solved enzyme:substrate Michaelis complex of ROC1 AzrC-MG (Figure 6; PDB-ID 9C0W) revealed that the distance of FMN N5 to the cationic imine was similar to the carbonyl carbon of orange I and acid red 88 in AzrC from *B.* sp. B29. This proximity suggested that reduction likely occurs at the imine of MG, resulting in the formation of LMG, for which a mechanism has been proposed. LCMS analysis supported this hypothesis as LMG was the main reaction product; we also noted the formation of desmethyl-LMG, but this formation was unexplained, potentially forming spontaneously.

In conclusion, we isolated and characterised the halophilic *Bacillus pacificus* ROC1 which was capable of aerobically degrading and detoxifying MG. We further identified the enzyme AzrC as capable of reducing MG to LMG, a process potentially involved during *in vivo* degradation. A potential mechanism for the reduction of MG by AzrC was suggested, supported by mass spectrometry analysis of the reaction products and the structure of the Michaelis complex, as determined by X-ray crystallography.

## Materials and methods

### Inoculum preparation

*Bacillus pacificus* ROC1 used in this work was isolated from the Kohat salt range, Kohat, Khyber Pakhtunkhwa, Pakistan. The whole ROC1 genome was submitted under the accession number SAMN33627856 to the NCBI.

### Media screen

To determine the ability of ROC1 to use MG as a sole carbon or nitrogen source, ROC1 was streaked onto three different minimal media. These media contained no carbon or nitrogen source (MM), contained glucose as a carbon source but no nitrogen source (MMG), or contained ammonium nitrate as a nitrogen source but no carbon source (MMN), and were composed as described previously, with the addition of 50 ppm MG (Fareed et al. 2017). Isolated bacteria was used to inoculate each medIa and the growth was monitored as described previously (Fareed et al. 2017).

### Biodegradation experiments

To monitor the degradation of MG, both free and immobilized cells were employed. Dye decolourisation experiments were performed in flasks containing different concentrations (0, 50, 100, 250, 400, 500, 750 and 1000 ppm) of MG. In all decolourisation experiments, percentage decolourisation was calculated using the formula:

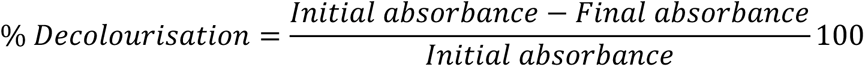

### Malachite green degradation by free cells

*E. coli* grown in LB media was supplemented with varying concentrations of MG, which was incubated at 37°C for 6 days. Every 24 hours, 3 mL of each sample was centrifuged at 8,000 rcf for 10 minutes, and the supernatant absorbance at 620 nm was measured, followed by calculation of percentage decolourisation using the previous equation, the pellet was used for mass determination. Free cell growth was monitored by OD_600_.

The CFU of free cells was measured every 24 hours. Five-fold serial dilutions of each sample were prepared and 20 mL of the 5^th^ dilution was spread onto LB agar, followed by incubation at 37°C overnight.

Free cell wet and dry biomass were also checked every 24 hours by weighing the cell pellet to determine the wet biomass, which was then dried at 90°C overnight using a heat block to determine the dry biomass.

### Effect of glucose and LB

For the 1,000 ppm MG samples, 0.5% (w/v) and 1% (w/v) glucose was added to media as an additional nutrient supply every 24 hours. Similarly, 1% (v/v) LB was added to the culture every 24 hours.

### Immobilisation of bacterial isolates

An agar entrapment method described previously was used (Fareed et al. 2022; Woodward 1988). Agar blocks containing ROC1 were prepared as described previously (Fareed et al. 2022).

### Malachite green degradation by immobilised cells

Three agar cubes containing ROC1 were added to LB media and varying MG concentrations were incubated with the cells for 6 days. Analysis was performed as for free cells.

### Bacteriotoxic effects

Bacteriotoxic effects of untreated and treated MG samples were assess by the agar diffusion methods with some modifications as described previously (Jamil et al. 2012; Phulpoto et al. 2018). *E. coli* and *S. aureus* were used, and were cultures in LB at 37°C at 220 rpm. Aliquots (OD_600_ equal to 0.5 MacFarland’s turbidity standard) were spread onto LB agar. Wells, 6 mm in diameter, were prepared and 50 μL sample from treated and untreated MG were loaded into the wells. Kanamycin (500 mg/mL) was used as a positive control. Plates were incubated at 37°C for 24 hours, and the clear zone around each well was measured.

### Phytotoxic effects

The commercially important tomato plant *Solanum lycopersicum* variety Riogrande was used for phytoxicity investigation of treated and untreated MG. Seeds (15) were sterilised with 1.2% (v/v) sodium hypochlorite, then were washed three times with water. Seeds were placed into Petri dishes containing filter paper moistened with treated or untreated MG solutions, followed by incubation at 22-24°C under light. Total seed germination was counted on days 3 and 5, the root and shoot length was measured, and percentage germination was calculated using the formula:

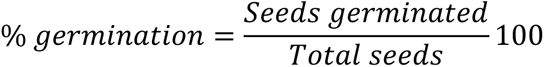

### Cell free extract enzyme assay

Reductase activity was determined by providing MG as a substrate and the CFE as an enzyme source. Reactions contained 20 mM Tris pH 7.4, 100 mM NADH, and CFE; MG decolourisation was monitored spectrophotometrically at 620 nm.

Laccase activity was determined by the ABTS method using 2,2’-azino-bis(3-ethylbenzothiazoline-6-sulfonic acid) (Roopa and Usha 2017). Reactions contained 0.5 mM ABTS, 0.1 M sodium acetate pH 4.5, and CFE. ABTS oxidation was monitored at 620 nm.

### Whole genome sequencing and analysis

#### DNA preparation and sequencing

ROC1 genomic DNA was extracted using the PureLink^TM^ genomic DNA mini kit. DNA was quantified by measuring absorbance at 260 nm using a NanoDrop®, followed by 1% (w/v) agarose gel electrophoresis. The ROC1 genome was sequenced by Illumina sequencing technology with 23X genome coverage.

#### Assembly and annotation

The genome was assembled using Geneious *de novo* assembler and gaps wre removed by rearrangement of contigs using Mauve multiple genome alignment software, the default progressive Mauve was used with the reference strain *Bacillus pacificus* NCCP 15909 (Accession number CP041979.1) (Darling et al. 2004). The sequenced genome was submitted to the NCBI under the accession number JARGCX000000000.1.

#### Phylogenetic analysis

The list of related species for ROC1 was obtained from LPSN (Parte et al. 2020). All accession numbers and genome sequences required were sourced from the NCBI. The list of all organisms used in phylogenetic analysis are displayed in Supporting Information 11. For phylogenetic tree construction, TYGS was used (Meier-Kolthoff and Goker 2019). Trees were constructed based on the whole genome of the selected organisms. The genome sequence data and related species accession numbers were uploaded to TYGS. Average nucleotide identity and AAI were calculated by the EDGAR platform, using all species included in the tree (Dieckmann et al. 2021).

#### Prediction of dye degrading genes

Two approaches were used to predict genes involved in MG degradation. First, RASTtk was used for annotation of ROC1, which annotated and divided the genome into sub-systems, from which the required genes were selected (Brettin et al. 2015). Second, the local BLASTp feature of BioEdit 7.2 was also used for identification of MG degrading genes. Protein sequences of enzymes involved in degradation from reference strains were sourced from IMG (Chen et al. 2023; Markowitz et al. 2006). These IMG sequences were used as a BLAST query against the ROC1 genome, using BLOSUM62 and an E-value 10.

#### Protein production and purification

MDF0734648.1 (AzrC), MDF0737692.1, and MDF0734684.1 were cloned into pET29-b(+) with a N-terminal hexahistidine tag and TEV cleavage site, under the control of a T7 promoter. The plasmids were transformed into *E. coli* BL21 (DE3) and transformants were cultured in LB containing 50 μg/mL kanamycin at 30°C at 180 rpm to OD_600_ 0.7-0.8, at which point gene expression was induced with 500 μM IPTG and 500 μM riboflavin, followed by incubation at 30°C at 180 rpm overnight. Cells were harvested by centrifugation at 4,730 rcf for 30 minutes at 4°C and were resuspended in 20 mM Tris, 200 mM NaCl, pH 8.0, followed by lysis via sonication and lysate clarification by centrifugation at 17,400 rcf for 60 minutes at 4°C.

Lysate was filtered through a 0.45 μm syringe filter and loaded onto a HisTrap^TM^ HP 5 mL column, followed by washing with 20 mM Tris, 200 mM NaCl, pH 8.0, 20 mM imidazole, and elution with 20 mM Tris, 200 mM NaCl, pH 8.0, 500 mM imidazole with a 0-100% B gradient over 20 CV. Fractions containing enzyme were concentrated and filtered through a 0.2 μm syringe filter, then loaded onto a HiLoad 26/600 Superdex® 200 pg SEC column. Protein purification was analysed by SDS-PAGE and Image Lab version 6.1 (Bio-Rad).

#### Azoreductase assay

All enzymes were buffer exchanged into 20 mM sodium phosphate, 200 mM NaCl, pH 5.0. Assays were performed using 50 μM enzyme, 2 mM NAD(P)H, 2 mM FMN, and 1 mM MG in different combinations, and absorbance at 620 nm was monitored.

#### Michaelis-Menten kinetics

Michaelis-Menten kinetics were determined using 50 μM AzrC, 2 mM NADPH, and MG concentrations of 20, 40, 80, 160, 320, 640, and 1280 μM, and absorbance at 620 nm was monitored. Data were processed using the Michaelis-Menten model – nonlinear regression in GraphPad Prism.

#### Reaction product determination

Reactions containing 50 μM AzrC, 3 mM NADH, and 1 mM MG were incubated at room temperature for two hours. The reaction was quenched with addition of 2X volume acetonitrile with 0.1% (v/v) formic acid. Quenched reactions were then filtered via centrifugation at 13,000 rcf for 15 minutes using an Amicon® Ultra Centrifugal Filter, 10 kDa MWCO. This was then diluted 1:100 in MQ with 0.1% (v/v) formic acid.

Reaction products were determined using LCMS using a gradient of 5-95% (v/v) acetonitrile with 0.1% (v/v) formic acid over 20 minutes with an ACQUITY^TM^ Premier BEH C18 1.7 μm VanGuard^TM^ FIT 2.1 x 50 mm Column attached to an UltiMate 3000 HPLC and a Q Exactive Plus mass spectrometer. 1 μL sample was run at 200 μL/min through the LC column, and was analysed by positive mode ESI with spray voltage 3.5 kV. Compounds were fragmented with 30 eV normalised collision energy.

#### Protein crystallisation

AzrC was purified as in section 4.11 with the addition of excess FMN to solution immediately prior to loading onto the SEC column.

AzrC (37 mg/mL) was incubated with 500 μM MG and 500 μM NAD^+^ before crystallisation. Crystals were grown at 19°C using 0.1 M bis-tris pH 5.5, 0.1 M sodium acetate, 20% (w/v) PEG 10K as mother liquor (ML) by hanging drop diffusion. AzrC was crystallised at a 1:1 protein:ML ratio. All crystals were manually looped and flash-cooled in liquid nitrogen.

#### X-ray crystallography data collection and processing

Diffraction data were collected on beamline MX2 at the Australian Synchrotron (Aragao et al. 2018). Data were collected over 360° with 0.1° rotation at a distance of 200 mm and 80% attenuation of the beam.

Data were indexed and integrated using XDSGUI, followed by resolution estimation and data truncation using AIMLESS as implemented in CCP4 (Agirre et al. 2023; Brehm et al. 2023; Evans and Murshudov 2013). The structure was solved by molecular replacement using Phaser MR as implemented in CCP4, using PDB entry 3w7a as the search model (Agirre et al. 2023; McCoy 2007; McCoy et al. 2007; Yu et al. 2014). MG was identified in difference density, and a Polder map was used to confirm MG coordinates (Liebschner et al. 2017). The model was iteratively refined and optimised using phenix.refine and Coot version 0.9.8.7 (Afonine et al. 2012; Emsley et al. 2010). The structure was validated using MolProbity. Phenix version 1.21.1 produced the refinement statistics (Table 2) (Afonine et al. 2010; Liebschner et al. 2019). The final structure was visualised in PyMOL version 3.0.3 (DeLano 2004; Schrodinger 2010). The MG environment and interactions were visualised using FLEV, as integrated into Coot, and PoseView (Emsley 2017; Stierand et al. 2006; Stierand and Rarey 2010).The structure factor CIF file was generated using pdb_extract and was validated using the wwPDB Validation System (Yang et al. 2004).

#### Statistical testing

All significance values were determined using a two-way ANOVA followed by a Tukey’s multiple comparisons test for when multiple datasets were being compared, or a Sidak’s multiple comparisons test for when two datasets were being compared.

### Accession codes

- AzrA – NCBI: MDF0737692.1
- AzrC – NCBI: MDF0734648.1
- YhdA – NCBI: MDF0734684.1

## Acknowledgements

This research was undertaken in part using the MX2 beamline at the Australian Synchrotron, which is part of ANSTO, and made use of the Australian Cancer Research Foundation (ACRF) detector.

