## Supporting Information for "Isolation, identification, and characterisation of the malachite green detoxifying bacterial strain *Bacillus pacificus* ROC1 and the azoreductase AzrC"

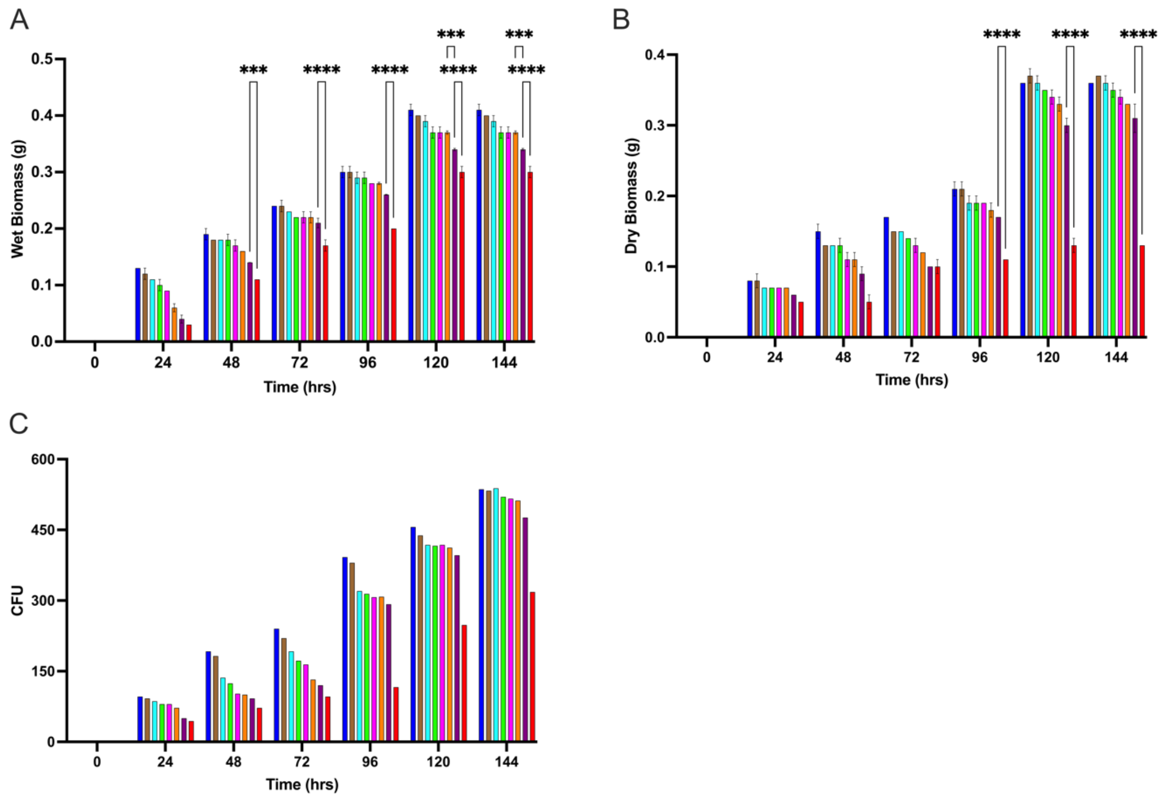


**Supporting Information 1. ROC1 growth with MG.** ROC1 growth over 144 hours with various MG concentrations: 0 ppm (blue), 50 ppm (brown), 100 ppm (cyan), 250 ppm (green), 400 ppm (magenta), 500 ppm (orange), 750 ppm (purple), and 1000 ppm (red). A – Wet biomass. B – Dry biomass. C – Colony forming units.

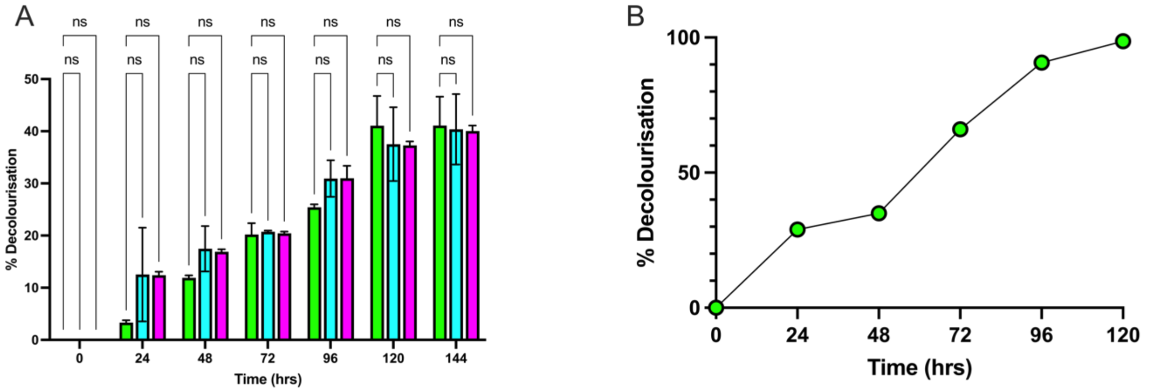


**Supporting Information 2. Modification of growth media.** Modification of media conditions for ROC1 decolourisation of 1,000 ppm MG. A – MG decolourisation over 144 hours with 0% glucose (green), 0.5% (w/v) glucose (cyan), and 1% (w/v) glucose (magenta). B – MG decolourisation over 120 hours with the addition of 1% (v/v) fresh LB media every 24 hours.

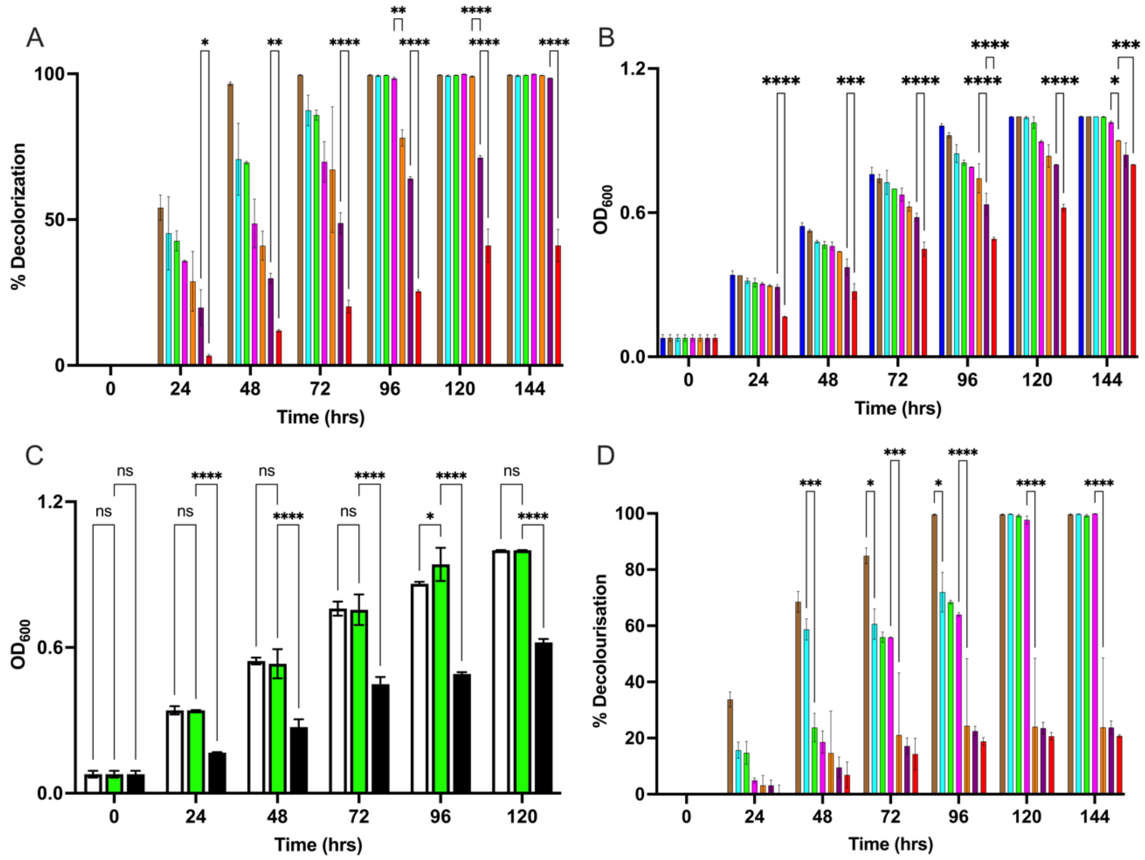


**Supporting Information 3. ROC1 decolourisation of MG.** A – Decolourisation of MG at 50 ppm (brown), 100 ppm (cyan), 250 ppm (green), 400 ppm (magenta), 500 ppm (orange), 750 ppm (purple), and 1000 ppm (red) measured at 620 nm over 144 hours by free ROC1 ±SD. Asterisks indicate significance (N=2). B – Optical density measured at 600 nm of ROC1 cultures grown over 144 hours in the presence of MG at 0 ppm (blue), 50 ppm (brown), 100 ppm (cyan), 250 ppm (green), 400 ppm (magenta), 500 ppm (orange), 750 ppm (purple), and 1000 ppm (red) ±SD. Asterisks indicate significance (N=2). C – Optical density of free ROC1 measured at 600 nm in the addition of 1% (v/v) media with 0 ppm (white) and 1000 ppm (green) MG, and with 1000 ppm without media addition (black) ±SD. Asterisks indicate significance (N=2). D - Decolourisation of MG at 50 ppm (brown), 100 ppm (cyan), 250 ppm (green), 400 ppm (magenta), 500 ppm (orange), 750 ppm (purple), and 1000 ppm (red) measured at 620 nm over 144 hours by immobilised ROC1 ±SD. Asterisks indicate significance (N=2).

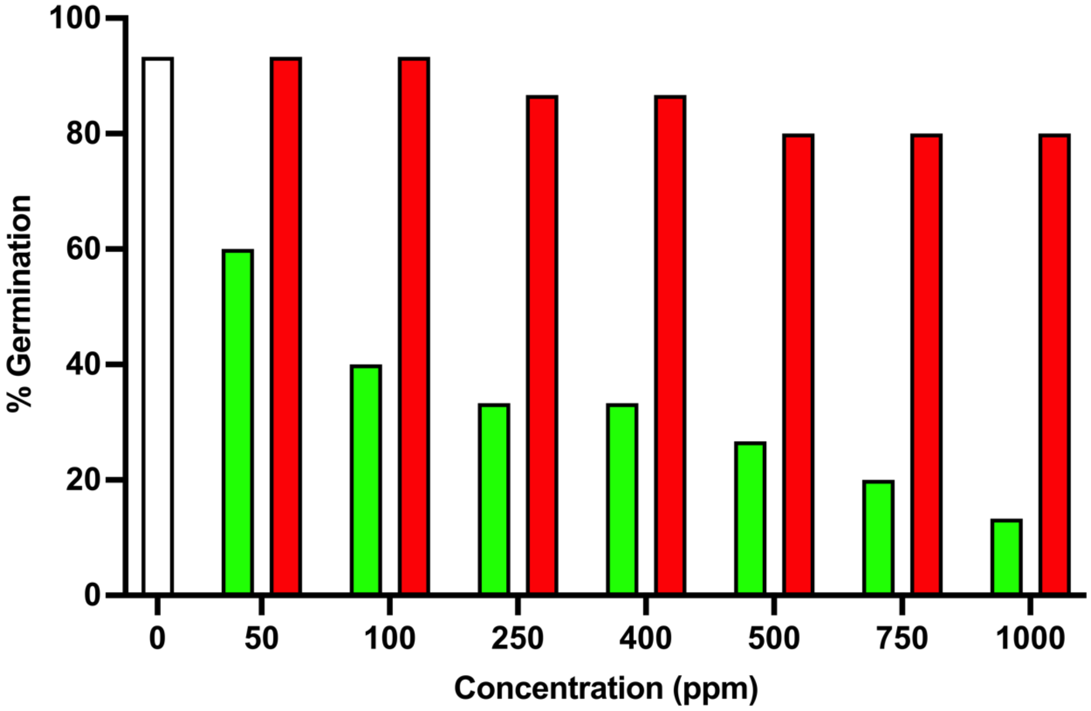


**Supporting Information 4. Seed germination after ROC1 treatment.** Percentage germination of *Solanum lycospersicum* seeds when incubated with water (white), untreated MG (green), and ROC1 treated MG (red).

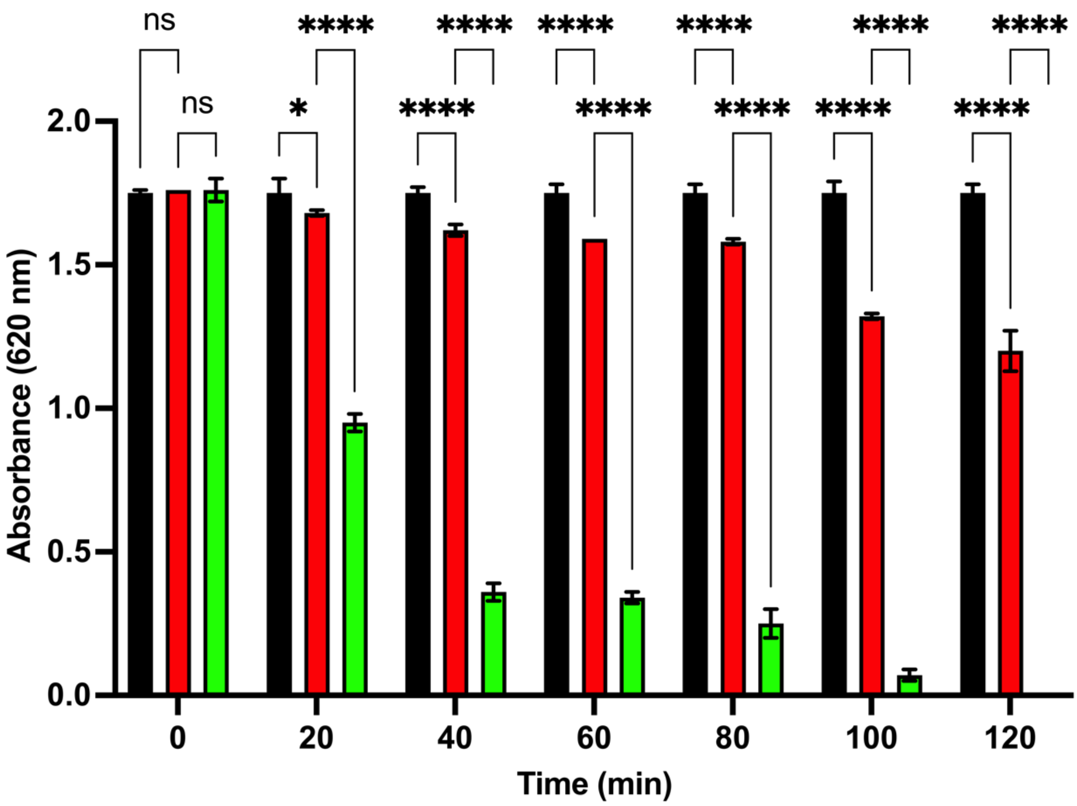


**Supporting Information 5. Reductase activity in the CFE.** MG absorbance (620 nm) measured over two hours in buffer (black), buffer with CFE (red), and buffer with CFE and NADH (green) ±SD. Asterisks indicate significance (N=3).

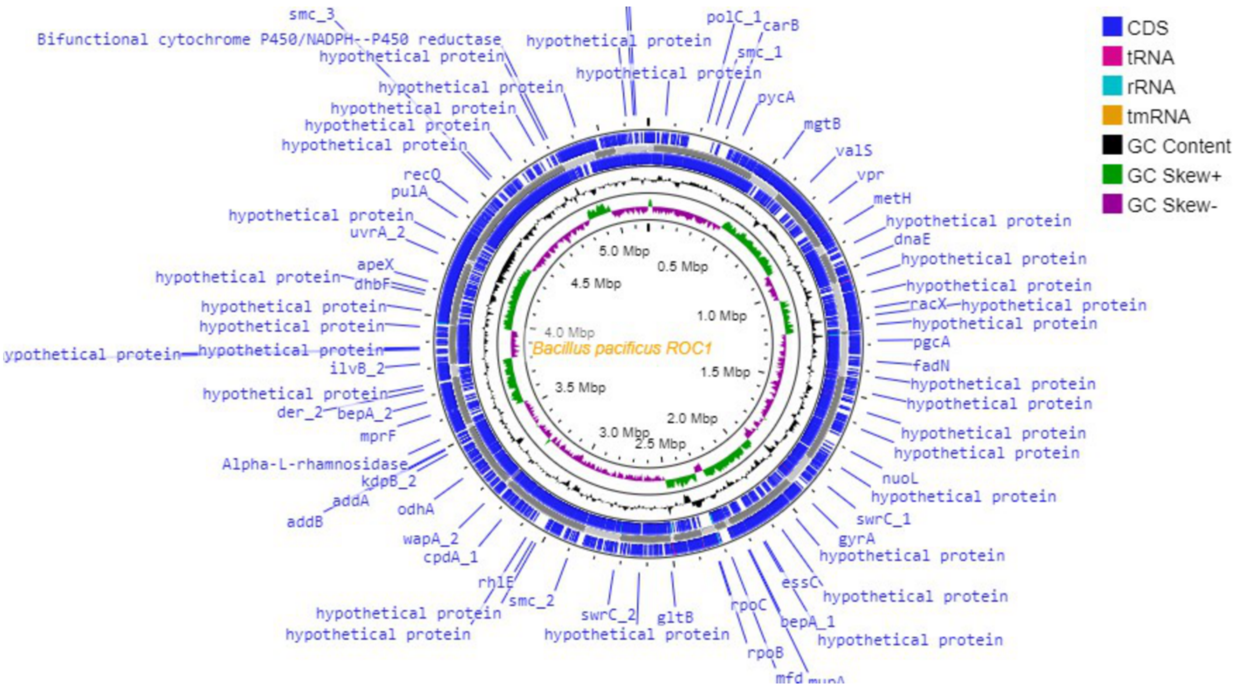

**Supporting Information 6. ROC1 Genome.** Circular genome of ROC1 generated using Proksee.

**Supporting Information 7.**

**Enzyme sequences.** Amino acid sequences of the three enzymes tested for their ability to decolourise MG, with a N-terminal TEV cleavage site (underlined) and hexa-histidine tag.

>MDF0734648.1

MGSSHHHHHHSSGENLYFQGMTKVLFITANPNSAEGSFGMAVGEAFIEAYKNEHP

QDEVVTIDLFNTTVPAIDADVFAAWGKFAAGEGFEALTEAQQQKVAAMNTNLETFM

HADRYVFVTPMWNFSYPPVVKAYLDNLAIAGKTFKYTENGPVGLLEGKKALHIQAT

GGVYSEGAYAAVDFGRNHLKTVLGFIGVNETEYIAVEGMNANPEKAQEIKEAAIANA

RELAKRF*

>MDF0737692.1

MGSSHHHHHHSSGENLYFQGMATVLFVKANNRPAEQAVSVKLYEAFLANYKEAHP

NDTVVELDLYKEELPYVGVDMINGTFKAGKGFDLTEEEAKAVAVADKYLNQFLEADK

VVFGFPLWNLTIPAVLHTYIDYLNRAGKTFKYTPEGPVGLIGDKKIALLNARGGVYSE

GPAAEVEMAVKYVASMMGFFGATNMETVVIEGHNQFPDKAEEIIAAGLEEAAKVAS

KF*

>MDF0734684.1

MGSSHHHHHHSSGENLYFQGMKLVVINGTPRKFGRTRVVAKYIADQFEGELFDLAV

EELPLYNGEESQRELEAVKKLKALVKNADGVVLCTPEYHNAMSGALKNSLDYLSSS

EFVHKPVALLAVAGGGKGGINALNSMRTVARGVYANAIPKQVVLDGLHVQDGELGE

DAKPLIHDLVKELKAYMGVYKEVKKQLGVE*

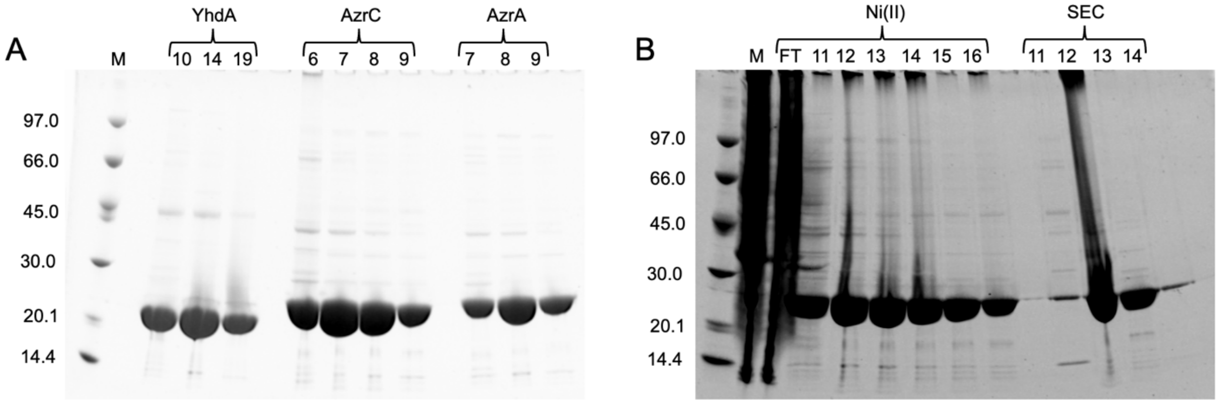


**Supporting Information 8. Azo-reductase purification.** SDS-PAGE analysis of azo-reductase purification. A – Elution fractions after Ni(II)-affinity chromatography for each protein. B – Elution fractions after Ni(II)-affinity chromatography (Ni(II)) and size exclusion chromatography (SEC) for AzrC purification.

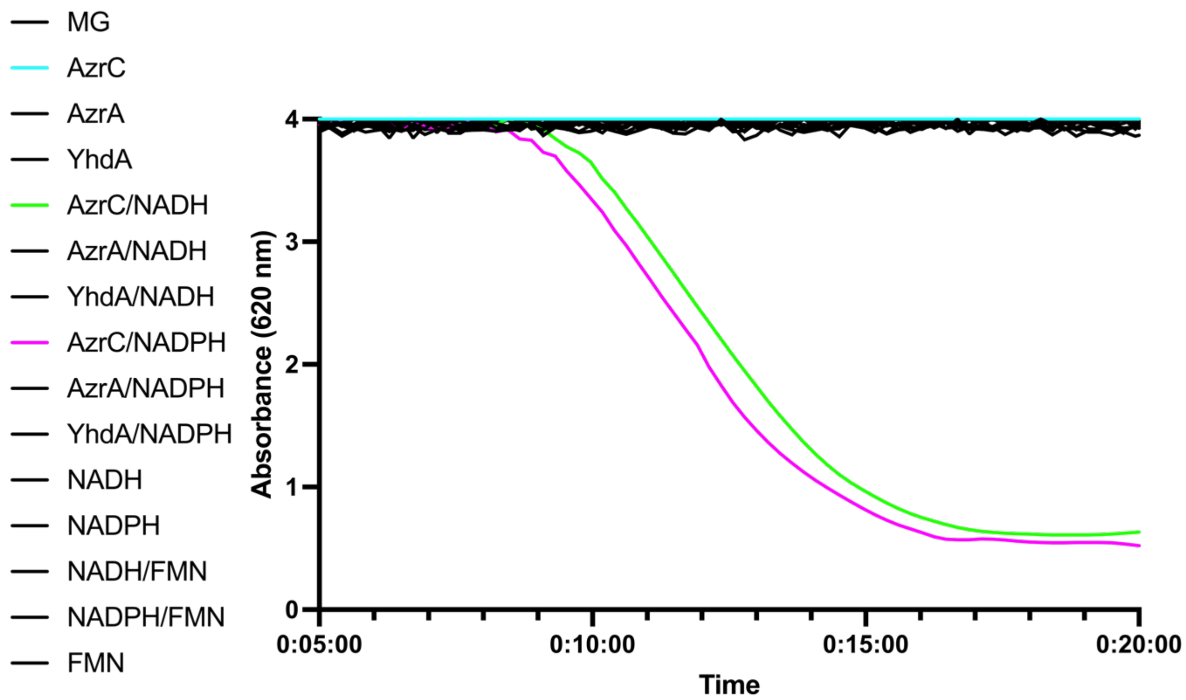


**Supporting Information 9. Initial MG decolourisation assay.** Absorbance of MG (620 nm) measured over 20 minutes. AzrC with NADH (green) and NADPH (magenta) was capable of MG deolourisation.

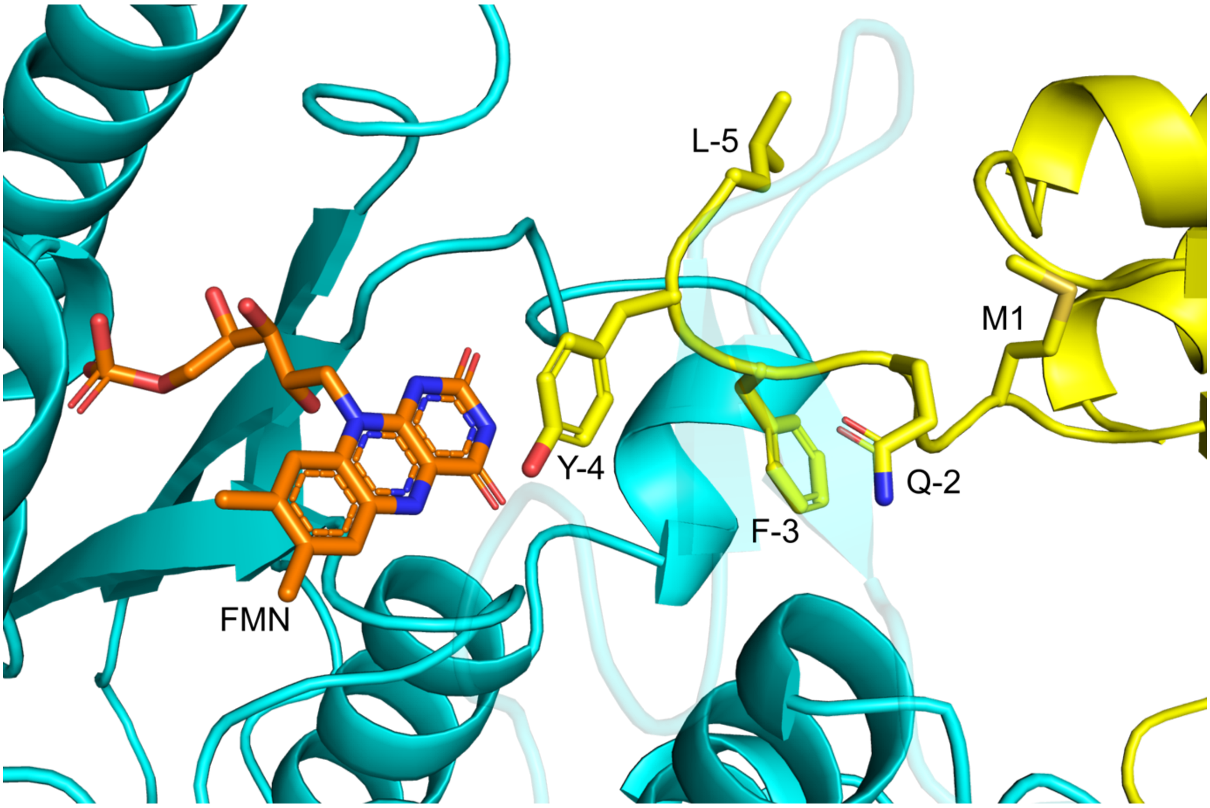


**Supporting Information 10. 9C0W symmetry mate.** The N-terminal sequence of a symmetry mate (yellow) occupies the active site of AzrC (cyan), preventing MG binding.

**Supporting Information 11. List of organisms.** The strains used in phylogenetic analysis of *Bacillus pacificus* ROC1 with the name, NCBI accession number, genome base pairs (bp), genome GC concent (GC), and the number of proteins encoded in the genome.

| **Name** | **Accession No.** | **bp** | **GC (%)** | **No. proteins** |
|---|---|---|---|---|
| *Bacillus cereus* A1 | NZ CP015727.1 | 5,352,307 | 35.28 | 5273 |
| *Bacillus thuringiensis* H3 | NZ CP052061.1 | 5,466,713 | 35.41 | 5480 |
| *Bacillus subtilis* N1142-3at | NZ CP064096.1 | 4,215,549 | 43.51 | 4227 |
| *Bacillus badius* NBPM-293 | NZ CP082363.1 | 3,868,812 | 44.18 | 3911 |
| *Bacillus tropicus* CK18 | NZ CP085399.1 | 5,237,233 | 35.48 | 5269 |
| *Bacillus anthracis* CMF9 | NZ CP085402.1 | 5,324,354 | 35.39 | 5336 |
| *Bacillus pacificus* ROC1 | JARGCX000000000.1 | 5,189,869 | 35.36 | 5254 |
| *Bacillus pacificus* NCCP 15909 | CP041979.1 | 5,126,903 | 35.61 | 5451 |
| *Bacillus cereus* ATCC 10987 | NC 003909.8 | 5,224,283 | 35.58 | 5276 |
| *Bacillus pacificus* anQ-h4 | NZ CP086328.1 | 5,252,926 | 35.53 | 5267 |
